# Differentially expressed growth factors and cytokines drive phenotypic changes in transmissible cancers

**DOI:** 10.1101/2024.11.06.622346

**Authors:** Kathryn G. Maskell, Anna Schönbichler, Andrew S. Flies, Amanda L. Patchett

## Abstract

The Tasmanian devil is threatened by two deadly transmissible Schwann cell cancers. A vaccine to protect Tasmanian devils from both devil facial tumour 1 (DFT1) and devil facial tumour 2 (DFT2), and improved understanding of the cancer cell biology, could support improved conservation actions. Previous transcriptomic analysis has implicated phenotypic cellular plasticity as a potential immune escape and survival mechanism of DFT1 cells. This phenotypic plasticity facilitates transition from a myelinating Schwann cell to a repair Schwann cell phenotype that exhibits mesenchymal characteristics. Here, we have identified cytokines and growth factors differentially expressed across DFT cell phenotypes and investigated their role in driving phenotypic plasticity and oncogenic properties of DFT cells. Our results show that NRG1, IL16, TGFβ1, TGFβ2 and PDGFAA/AB proteins have significant and distinct effects on the proliferation rate, migratory capacity and/or morphology of DFT cells. Specifically, PDGFR signalling, induced by PDGFAA/AB, was a strong enhancer of cell proliferation and migration, while TGFβ1 and TGFβ2 induced epithelial-mesenchymal transition (EMT)-like changes, inhibited proliferation and increased migratory capacity. These findings suggest complex interactions between cytokine signalling, phenotypic plasticity, growth and survival of DFTs. Signalling pathways implicated in the propagation of DFT are potential targets for therapeutic intervention and vaccine development for Tasmanian devil conservation.

## Introduction

The Tasmanian devil (*Sarcophilus harrisii*), a carnivorous marsupial native to the Australian island state of Tasmania, is threatened by two deadly transmissible cancers: Devil Facial Tumour 1 (DFT1), discovered in 1996, and Devil Facial Tumour 2 (DFT2), discovered in 2014. Both cancers share gross similarities and a common origin from Schwann cells, a plastic cell type capable of changing phenotypically to control myelination and repair functions in the peripheral nervous system (1–3). Despite gross similarities, genetic karyotyping, major histocompatibility class 1 (MHCI) typing, and microsatellite analyses have revealed that DFT1 and DFT2 are distinct, and arose independently in different Tasmanian devils (4–6). The spread of DFT1 across the majority of mainland Tasmania has resulted in regional population declines of over 82%, with severe impacts on local ecosystems (7, 8). In contrast, DFT2 is currently detected only in the southeast region of Tasmania (9). The potential impact of DFT2 on the already diminished populations, should it spread further across Tasmania, remains unknown. As such, a vaccine to protect Tasmanian devils from both DFT1 and DFT2 is a conservation priority (10).

DFTs represent an intriguing paradigm, whereby the tumour cell itself is the infectious pathogen and is transferred as an allograft via biting behaviours (4, 11). Typically, immune defences targeting non-self, allogeneic cells should be activated against transmitted tumour cells. However, DFT allografts fail to evoke these protective responses, developing into malignant tumours in new hosts (12). Initial studies into the lack of immune-mediated DFT rejection found that DFT1 cells exhibit epigenetic downregulation of essential components of the MHCI antigen processing pathway, including β2-Microglobulin (β2M) (13). The downregulation of these genes prevents MHCI display at the DFT1 cell surface, hindering T-cell recognition. Efforts to generate a prophylactic DFT1 vaccine have focussed on reversing this MHCI downregulation by treating DFT1 cells with interferon-gamma (IFNγ) to upregulate MHCI display for vaccination (14–18). Although this vaccine design was unsuccessful at preventing DFT1 infection, 95% of immunised devils exhibited an increase in antibody-mediated tumour reactivity, and lymphocyte infiltration into the tumours was observed in immunised devils (14, 19). This suggests that vaccine-mediated DFT1-specific immune responses were activated but ultimately circumvented by a secondary immune escape mechanism. Similarly, studies have revealed that DFT2 can persist despite retaining MHCI expression (5, 20). This supports the theory that alternative mechanisms of immune evasion combine with MHCI aberrations to prevent immune-mediated DFT rejection in Tasmanian devils.

To better understand the mechanisms facilitating DFT immune escape in Tasmanian devils, Patchett *et al*., 2021 conducted transcriptomic analyses on DFT1 tumours established in both immunised and non-immunised Tasmanian devils (21). When an unvaccinated Tasmanian devil was challenged with DFT1, the resulting tumour exhibited a semi-differentiated ‘myelinating’ Schwann cell phenotype. In contrast, DFT1 tumours that developed in vaccinated Tasmanian devils displayed a de-differentiated ‘mesenchymal’ phenotype. This phenotype was characterized by the upregulation of genes associated with epithelial-to-mesenchymal transition (EMT) and stemness, and the downregulation of myelin genes, such as myelin basic protein (MBP) and periaxin (PXN). The mesenchymal DFT phenotype was also accompanied by increased immune infiltration, highlighting that in the presence of immune cells and potential tumour recognition, this phenotype may facilitate immune escape. Indeed, DFT2, which is seemingly more immunogenic due to its sustained MHCI expression, naturally adopts a mesenchymal phenotype, remaining capable of transmission in this form. An important marker of the DFT2 mesenchymal phenotype is platelet-derived growth factor receptor alpha (PDGFRA) expression (3, 9), a known driver of EMT in human cancers (22).

An inherent plasticity of DFT cells in response to a changed immune environment could enable DFT survival in the presence of tumour-specific immune responses. Indeed, EMT-like changes are associated with altered immunogenicity in human cancers (23–26). Here, we have identified drivers of mesenchymal plasticity in DFT cancers and demonstrated significant effects on the oncogenic properties of DFT cells. Our findings have revealed novel survival mechanisms in DFT cancers and can be applied for future development of prophylactic and therapeutic interventions against DFTs in Tasmanian devils.

## Methods

### Tasmanian Devil proteins of interest

Predicted Tasmanian devil coding sequences for the proteins of interest (NRG1, IL16, TGFB1/2 and PDGFAA/AB) were acquired from the National Centre for Biotechnology Information (NCBI) reference assembly mSarHar 1.11. Each sequence was aligned against its respective human homologue, and if the sequence was conserved across species, the human protein was purchased commercially (TGFβ1/2, PDGFAA/AB). Where the sequence was not conserved (i.e. NRG1 and IL16), fluorescent, devil-specific recombinant proteins were developed using established methods (27, 28).

### Cell culture

DFT1 cell lines, C5065 (RRID: CVCL_LB79) and 1426 (RRID: CVCL_LB76) were established by A-M Pearse and K. Swift of the Tasmanian Department of Primary Industries, Parks, Water and Environment (DPIPWE). The lines originated from DFT1 tumour biopsies obtained under the approval of the Animal Ethics Committee of the Tasmanian Parks and Wildlife Service (permit numbers 33/2004-5 and 32/2005-6). The DFT2 cell line, Jarvis (JV) (RRID: CVCL_A1TN), was established by A. Kreiss and R. Pye of Menzies Institute for Medical Research, University of Tasmania (5). The cells originated from tumour biopsies obtained under the approval of the University of Tasmania Animal Ethics Committee (permit number A0012513). For all experiments, cell stocks thawed from –80 °C were maintained at 37 °C with 5% CO_2_ in RPMI medium (11875093, GIBCO, USA) supplemented with 10% heat-inactivated foetal calf serum (A31605-01, GIBCO, USA) and 1% Antibiotic Antimycotic (15240096, Thermo Fisher Scientific, USA) (CRF10).

### Bioluminescence imaging (BLI)-based proliferation assay

C5065 and JV cell lines previously stably transfected with a firefly luciferase encoding plasmid (pAF112; Addgene # 135920) were cultured in CRF10 with the selection reagent, hygromycin (H0654, Sigma Aldrich, USA), at 200 µg/mL (27). Each cell line was aliquoted into the appropriate wells of a 96-well, white wall, glass bottom plate (164590, Thermo Fisher Scientific, USA) at 5000 cells per well, and allowed to adhere at 37 °C 5% CO_2_ for 2 hours. VivoGlo D-Luciferin (P1041, Promega, USA) was added to a final concentration of 75 µg/mL. Treatment proteins were serially diluted in triplicate across wells by a factor of 10 from 25 to 0.025 ng/mL. The cells in maximum lysis control wells were lysed by adding 200 µL of sterile Milli-Q® water (Merck Millipore, USA) to the cells. The plates were incubated at 37 °C and 5% CO_2_ and bioluminescence was measured on the Tecan Spark® Microplate Reader (Tecan, USA) at 0 hours and every 24 hours for 7 days (inclusive). The data were analysed by 2-way ANOVA on GraphPad Prism version 9.0.0 (GraphPad Software, USA) with Dunnett’s multiple comparison test.

### Cell viability assay

To determine the half-maximal inhibitory concentration (IC50) of the TGFβ receptor I (TβRI) inhibitor Galunisertib (LY2157299, Selleckchem, USA) on DFT cell lines, cell viability assays were performed. C5065 and JV cell lines were seeded in 96-well flat-bottom plates (3598, Corning, USA) at a density of 20,000 cells (C5065) or 10,000 cells (JV) per well. Cells were then treated in triplicate with serial dilutions of Galunisertib or with 20% dimethyl sulfoxide (DMSO; D8418, Sigma Aldrich, USA) as a positive control for inhibition of cell proliferation. Cell viability was measured after 72 hours of incubation using a Cell Counting Kit-8 (CCK-8; 96992, Sigma-Aldrich, USA). Measurements were taken using a Tecan Spark® Microplate Reader (Tecan, USA) and IC50 values were calculated through non-linear regression analysis.

### Scratch-plate migration assay

C5065 and JV cells were plated at 1.65x10^5^ cells per well in a 24-well plate (3526, Corning, USA) and were allowed to adhere for 1 hour before the treatment proteins were added at 50 ng/mL (IL16, TGFβ1/2 and PDGFAA/AB) or 5 ng/mL (NRG1). Cells were incubated at 37 °C and 5% CO_2_ for 48 hours. Matrigel® basement membrane (CLS356234, Corning, USA) was diluted to 0.3 mg/mL in cold CRF10 and 20 µL was added to the appropriate wells of a cold Incucyte® Imagelock 96-well plate ( BA-04856, Sartorius, USA). The plate was incubated at 37 °C and 5% CO_2_ for 30 minutes. Pre-treated cells were plated in triplicate onto the prepared Incucyte® Imagelock 96-well plate at 5x10^4^ cells/well in 50 µL CRF10. The plate was incubated for 1 hour at 37 °C and 5% CO_2_ before treatment protein was added at 50 ng/mL (IL16, TGFβ1/2 and PDGFAA/AB) or 5 ng/mL (NRG1). Untreated and no-Matrigel controls were included in triplicate. Cells were incubated for a further 24 hours at 37 °C and 5% CO_2_. After 24 hours, the cells in the wells were ‘wounded’ using the Incucyte® Woundmaker tool (BA-04858, Satorius, USA) per the manufacturer’s instructions. Wells were rinsed with PBS and protein of interest was added to the scratched wells at 50 ng/mL (IL16, TGFβ1/2 and PDGFAA/AB) or 5 ng/mL (NRG1) and the plate was transferred to the Incucyte® live cell analyser. Each well was imaged every 2 hours on the 10X objective. A threshold for percent wound confluency was chosen for each cell line based on the minimum confluency reached by each treatment condition. The time point at which each replicate crossed the threshold was determined by fitting an exponential curve to the data and interpolating the time the sample crossed the threshold. Data was analysed by one-way ANOVA followed by Dunnett’s multiple comparisons test using GraphPad Prism version 9.0.0 (GraphPad Software, USA).

### Tasmanian devil peripheral blood separation

Tasmanian devil peripheral blood was collected from three Tasmanian devils into lithium-heparin tubes as previously described (29) (Ethics approval numbers 23230 and 26159, UTAS Animal Ethics Committee). Blood samples were diluted 1:1 with cold CRF0 (RPMI supplemented with 1% antibiotic-antimycotic and 1% GlutaMAX (35050061, Thermo Fisher Scientific, USA)) and peripheral blood mononuclear cells (PBMCs) were separated by density gradient centrifugation on Histopaque ® (10771, Sigma Aldrich, USA) as per the manufacturer’s instructions. The PBMC pellet was resuspended in CRF10 and quantitated by trypan blue (15250061, Invitrogen, USA) exclusion.

### Bioluminescence Imaging (BLI)-based ATP killing assay

C5065 and JV cell lines previously transfected with a firefly luciferase encoding plasmid (pAF112; Addgene # 135920) were cultured in CRF10 with the selection reagent, hygromycin, at 200 µg/mL (27). Cell lines were plated at 2000 cells per well in CRF10 and D-luciferin at a final concentration of 75 µg/mL. After 2 hours incubation at 37°C 5% CO_2_, treatment proteins (TGFβ1/2 and PDGFAB) were added in duplicate at a final concentration of 50 ng/mL. Untreated and maximum lysis controls were included by adding CRF10 or sterile Milli-Q® water respectively. The plates were incubated at 37°C 5% CO_2_ for 24 hours. PBMCs were added in duplicate to the treated cells at ratios of PBMC:target cell of 12:1, 25:1, 50:1 and 100:1. A no-PBMC control was included for each treatment condition (0:1 ratio). Recombinant Tasmanian devil interleukin-2 (IL2) (made by A Flies, Menzies Institute for medical research) was added to the co-culture at a final concentration of 50 ng/mL. Control wells without IL2 were included in duplicate. Plates were incubated at 37°C and 5% CO_2_ and luminescence (produced by the decomposition of D-luciferin by luciferase) were measured by the Tecan Spark® Microplate Reader at 4 hrs and 18 hrs. Duplicate wells were averaged and relative DFT death was estimated based on the luminescence count in the no-PBMC control well of each treatment condition divided by the luminescence count in the experimental well of the same treatment condition. Relative DFT death across treatment conditions was analysed by one-way ANOVA, comparing the main treatment effect to the no treatment control. Dunnett’s multiple comparisons test was used to compare treatment at individual cell ratios.

### Actin Staining

Sterile 12 mm coverslips (G401-12, ProSciTech, AUS) were added to a 24-well plate, treated with Poly-L-Lysine (P4707, Sigma Aldrich, USA) for 10 minutes, and UV treated for 30 minutes. C5065, 1426 and JV cell lines were plated at a density of 1.75x10^5^ cells per coverslip for 24-hour samples or 1.5x10^5^ cells per coverslip for 48 hour samples in CRF10. Plates were incubated for 2 hours at 37°C and 5% CO_2_ prior to the addition of treatment proteins at a final concentration of 50 ng/mL. After 24 or 48 hours, coverslips were washed with PBS (18912094, Thermo Fisher Scientific, USA) to remove treatment. Cells were fixed with 4% paraformaldehyde (PFA) for 20 minutes, rinsed with PBS, and permeabilised with 0.3% Triton X-100 (T9284, Sigma Aldrich, USA) for 10 minutes. Blocking buffer (0.3 M glycine, 1% BSA and 0.1% sodium azide) was added for 20 minutes before the cells were stained with Phalloidin-iFluor 488 (ab176753, Abcam, UK) for 20 minutes. Finally, the coverslips were rinsed in PBS and stained with DAPI (D9542 , Sigma Aldrich, USA) at 200 ug/ml for 10 minutes. Coverslips were mounted onto microscope slides in Fluorescence Mounting Medium (S3023, Dako Agilent, USA) and imaged using the Olympus BX50.

## Results

### Signalling proteins expressed in the DFT microenvironment alter DFT cell proliferation

DFT cancers exist in ‘myelinating’ (semi-differentiated) and ‘mesenchymal’ (dedifferentiated) phenotypes, which could be linked to key tumorigenic processes such as proliferation, migration, invasion and immune evasion (3, 30). To explore how these phenotypes influence tumorigenic functions, we identified potential drivers of DFT phenotypic plasticity using previously published transcriptomic data (3, 21). We chose five proteins of interest (POIs) and a subset of their receptors that were differentially expressed across varying DFT phenotypes and had previously been identified as having roles in driving human cancers, such as CD9 and its secondary receptor IL16 (Figure 1). We then treated DFT cells with serially diluted POIs and observed their effect on cell proliferation using relative light units (RLU) produced by bioluminescent cells. We used the cell lines C5065 and Jarvis (JV), which will henceforth be referred to as DFT1 and DFT2 cell lines, respectively. Our data indicated notable differences in proliferation rates of DFT1 and DFT2 cell lines, both at baseline (Supplementary figure 1) and after treatment with POIs (Figure 2). Specifically, the proliferation rate of DFT1 was significantly increased by PDGFAB (p < 0.0001) but inhibited by TGFβ1 (p < 0.0001) and TGFβ2 (p < 0.0001) (Figure 2B, E, and F). Conversely, treatment with NRG1, IL16, and PDGFAA did not significantly affect the proliferation rate of DFT1 cells (Figure 2A, C, and D). The proliferation of DFT2 cells was significantly inhibited by TGFβ1 (p < 0.0001) and TGFβ2 (p < 0.0001) (Figure 2H, K). However, NRG1, IL16, PDGFAA, and PDGFAB had no significant effect on the proliferation rate of DFT2 cells (Figure 2G, I, J, and L). These findings suggest that proliferation rate in DFT cancers is influenced by POIs associated with varied DFT phenotypes.

**Fig. 1.**
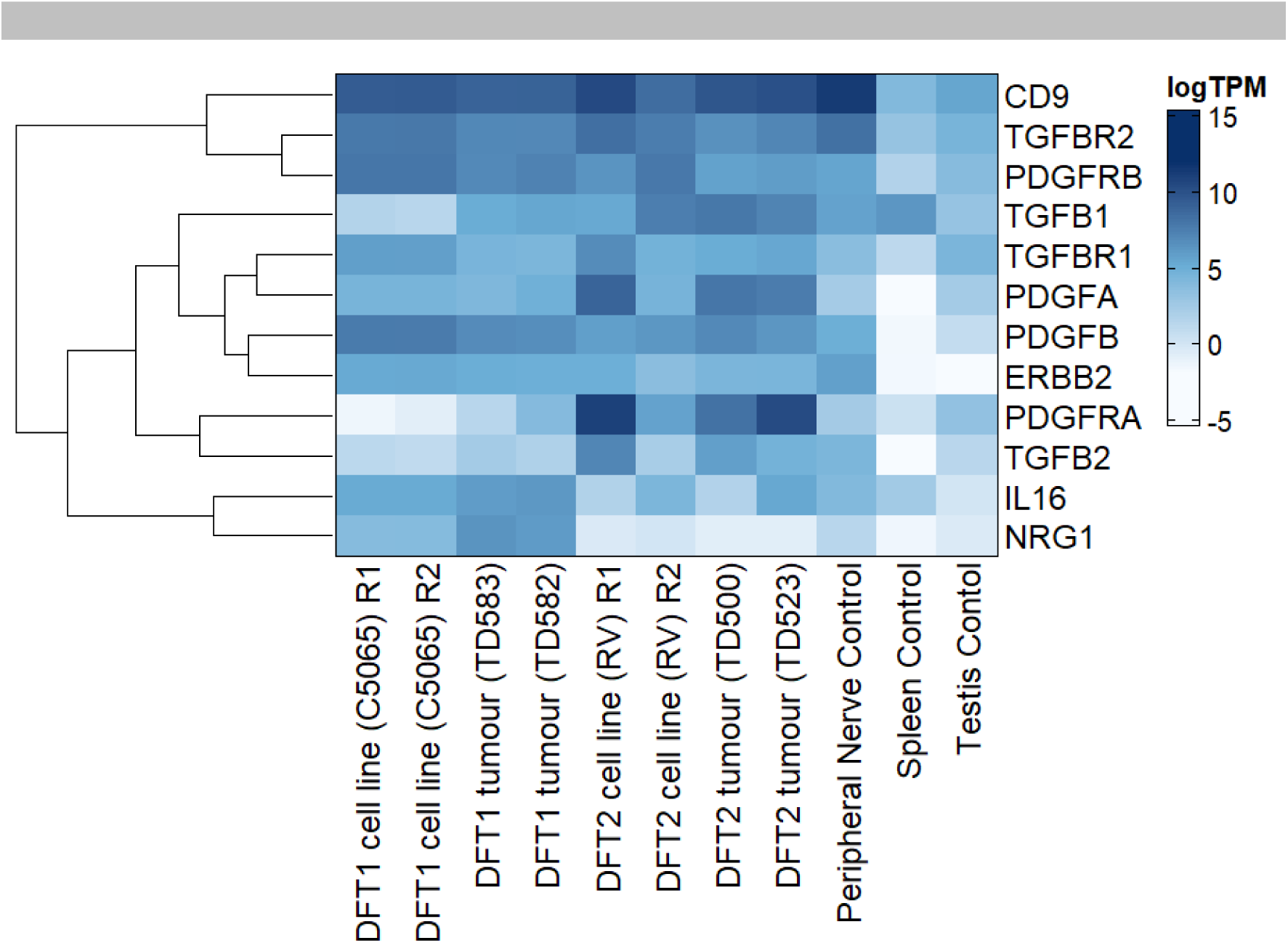
Heatmap depicting relative transcript abundance of ligand-receptor gene pairs in DFT cell lines and tumours. Devil peripheral nerve, testis and spleen are included as controls. Transcript abundance is represented by log-transcripts per million (TPM) and visualised from low to high on a gradient of white to navy blue. Gene expression is clustered by Euclidean distance. Transcript abundance data was obtained from Patchett et al (3). Heatmap was made with ComplexHeatmap v2.20.0 in R 4.4.0

**Fig. 2.**
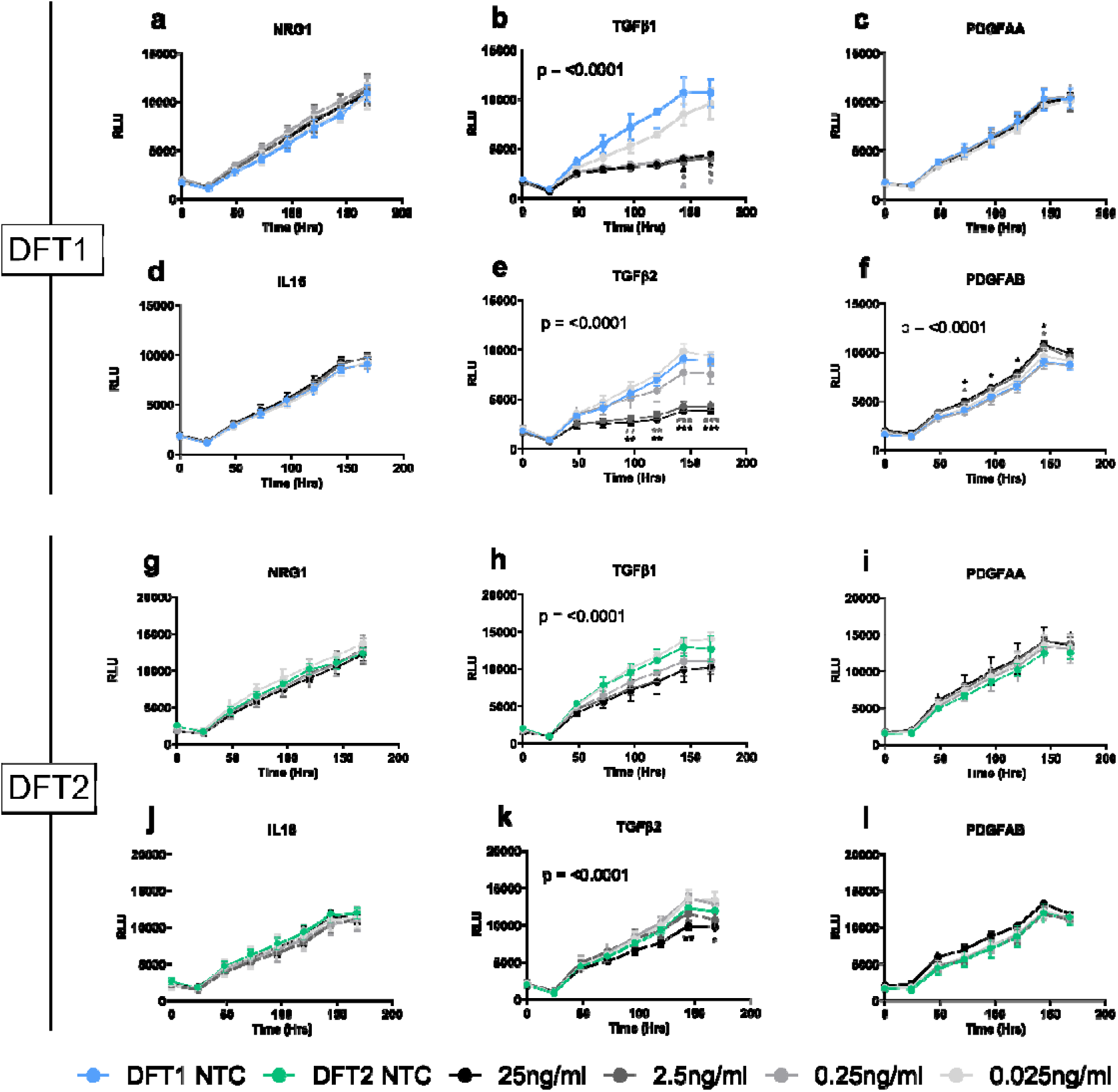
Bioluminescent imaging (BLI) – based proliferation assay. DFT1 and DFT2 cells were stably transfected with a firefly luciferase encoding plasmid, allowing the cells to emit bioluminescence in the presence of adenosine triphosphate and exogenous luciferin. (A-F) DFT1 cells and (G-L) DFT2 cells were treated with serially diluted proteins of interest (NRG1, IL16, TGFB1/2 and PDGFAA/AB) for 168 hours. Relative light units (RLU) were measured every 24 hours as a proxy of live cell number. Data for each treatment concentration were plotted as RLU over time (hours). Statistical significance compared to the no treatment control (NTC; 0 ng/mL) was assessed using a two-way ANOVA with Dunnett’s multiple comparisons test. P-values on graphs represent the significance of the overall interaction effect between time and treatment concentration. Statistical significance of multiple comparisons was set at p<0.05 and is indicated by asterisks: where: * =P<0.05; **=P<0.01; ***=P<0.001; and ****=P<0.0001. Asterisks are colour co-ordinated with the respective treatment. Each point represents the mean of data in triplicate with error bars representing the standard deviation. Graphs were created using GraphPad Prism version 9.0.0

### Dose-dependent effects of PDGFAA and PDGFAB on DFT cell proliferation

DFT cells express heightened levels of ERBB and PDGF receptors and their ligands NRG1, PDGFAA, and PDGFAB (Figure 1)(21). Considering the minor or negligible changes in DFT cell proliferation after the addition of low concentrations of these ligands (Figure 2), we opted to repeat our proliferation assays using higher concentrations. Although DFT1 cells are driven by ERBB3 signalling (31), the NRG1 isotype used in this study failed to stimulate significant proliferation of both DFT1 and DFT2 cells at high concentrations (Figure 3A, D; Supplementary figure 2, 3). In contrast, PDGFAA and PDGFAB significantly increased the proliferation of both cell lines (p < 0.0001) (Figure 3B-C, E-F; Supplementary figure 2, 3). This suggests that PDGF receptor signalling is active in both DFT1 and DFT2 and could play a primary role in promoting tumour growth.

**Fig. 3.**
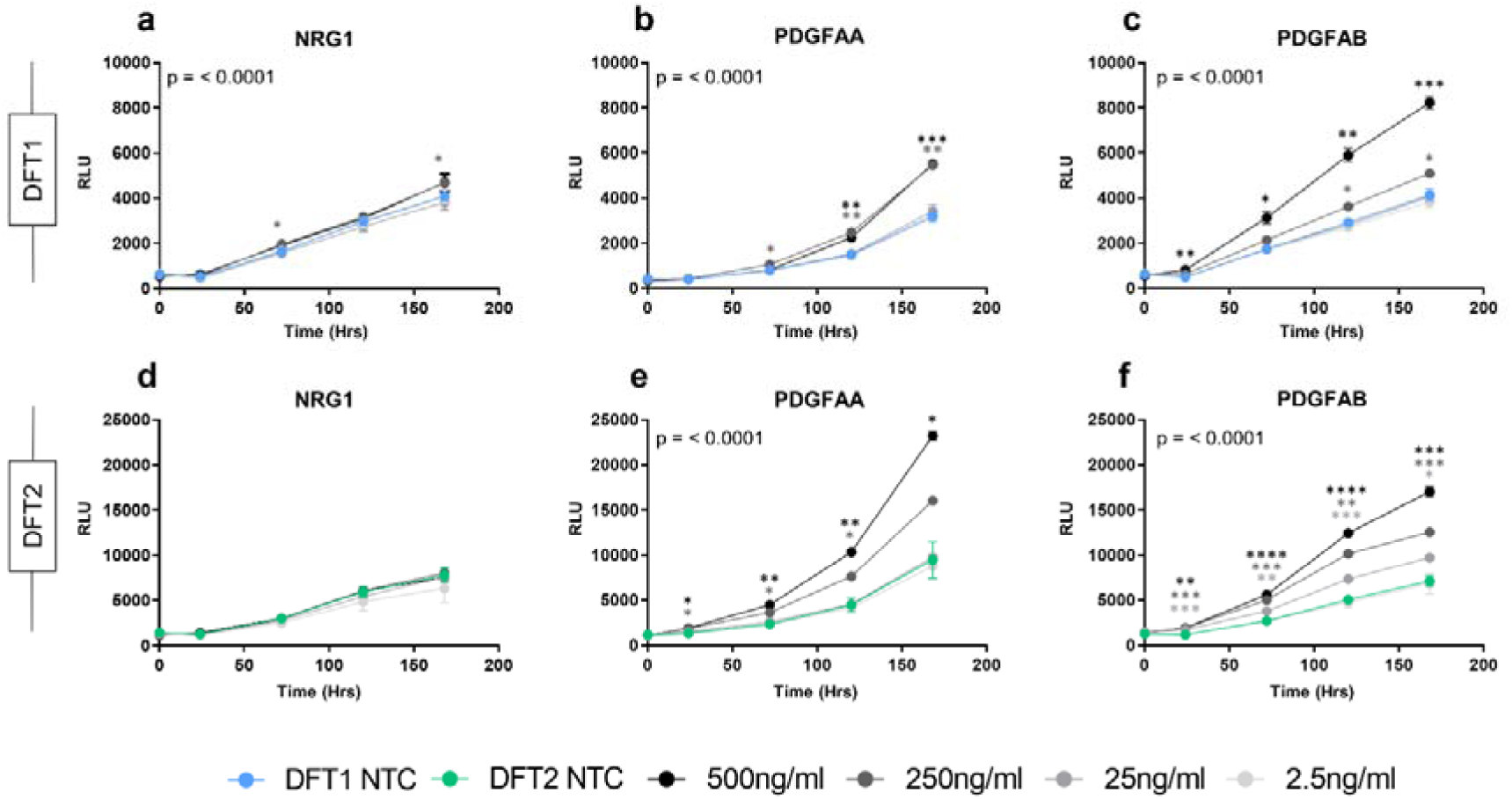
Bioluminescence imaging-based proliferation assay with high POI concentrations. (A-C) DFT1 cells and (D-F) DFT2 cells were treated with NRG1, PDGFAA and PDGFAB for 168 hours. Relative luminescence was measured every 24 hours as a proxy of cell number. Data for each treatment concentration were plotted as relative light units (RLU) over time. Statistical significance compared to the no treatment control (NTC; 0 ng/mL) was measured using a two-way ANOVA with Dunnett’s multiple comparisons test. P-values on graphs represent the significance of the overall interaction effect between time and treatment concentration. Statistical significance of multiple comparisons was set at p<0.05 and is indicated by asterisks: * =P<0.05; **=P<0.01; ***=P<0.001; and ****=P<0.0001. Asterisks are colour co-ordinated with the respective treatment. Each point represents the mean of data in triplicate with error bars representing the standard deviation. Graphs were created using GraphPad Prism version 9.0.0

### Signalling proteins expressed in the DFT microenvironment affect migratory capacity

Mesenchymal phenotypes in human cancers are associated with regulation of cell migration and metastasis (32–34). To determine whether POIs associated with DFT1 and DFT2 phenotypes impact the migratory potential of DFT cells, we performed scratch plate cell migration assays. For statistical analyses, confluency thresholds were applied to each cell line based on the minimum confluency reached by all treatment conditions. The wound confluence in the DFT2 cell line was significantly higher than the DFT1 cell line at every time point after 4 hours until the cessation of measurements (p=<0.0001) (Supplementary figure 4).

DFT1 cells treated with NRG1, IL16, PDGFAA and PDGFAB reached wound confluency significantly faster than the no treatment control (P= 0.0002, 0.002, 0.004 and 0.0002, respectively) (Figure 4A-B). Comparatively, there was no statistically significant difference in the time taken for TGFβ1 and TGFβ2 treated cells to reach this threshold, although an inhibitory trend was observed at later time points (> 48 hours). The migratory capacity of DFT2 cells was significantly enhanced by treatment with NRG1, IL16, TGFβ1, TGFβ2 and PDGFAB (P= 0.0002, 0.0001, 0.02, 0.0002 and <0.0001, respectively) (Figure 4C-D). PDGFAA was the only POI to not significantly impact DFT2 migration. These findings showed that proteins associated with DFT phenotypes can alter the migration rate of DFT cells.

**Fig. 4.**
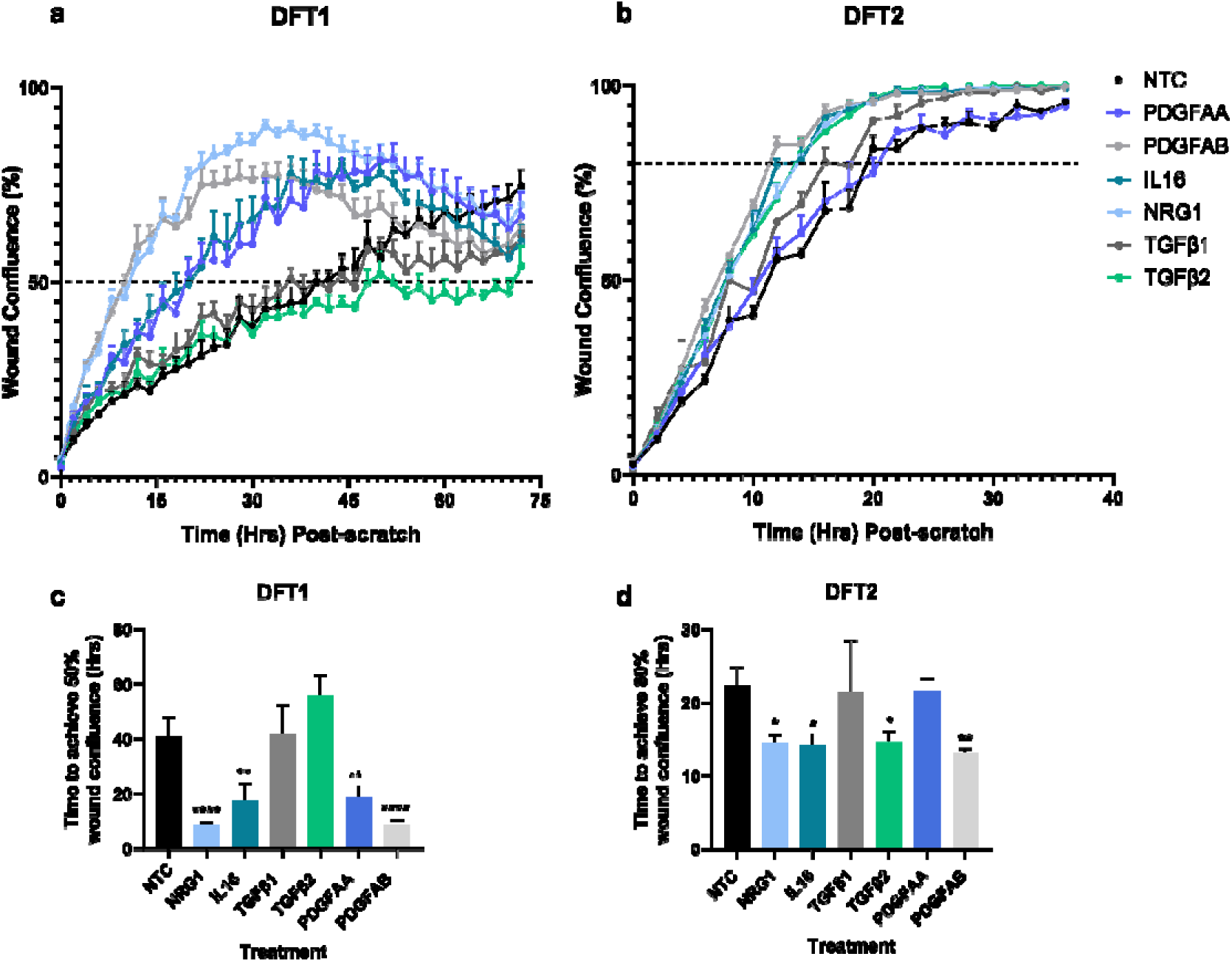
Scratch-plate migration assay. (A-B) DFT1 and (C-D) DFT2 cell lines were pre- incubated with NRG1, IL16, TGFβ1, TGFβ2, PDGFAA or PDGFAB at 0.5 ng/ml (NRG1) or 50 ng/mL for 72 hours on 0.3 mg/mL Matrigel. A 700-800 μm scratch wound was created down the centre of the well and a second dose of treatments was added. Images were taken every 2 hours. (A, C) Each point represents mean wound confluency with upward error bars showing standard deviation. The dashed lines at 40% and 80% on (A) DFT1 and (C) DFT2 graphs represent the confluency threshold value chosen for statistical analyses. (B, D) The time point at which each (B) DFT1 and (D) DFT2 replicate crossed the confluency threshold value was determined by fitting an exponential plateau curve to the confluency over time curves and interpolating the time the sample crossed the threshold from the equation of the line. The time point is represented here by mean with standard deviation. Statistical significance relative to the no treatment controls (NTC) was measured by one-way ANOVA followed by Dunnett’s multiple comparisons test. Statistical significance is illustrated as: *P<0.01; **P<0.01; ***P<0.001; ****P<0.0001. Graphs were created using GraphPad Prism version 9.0.0

### Treatment with TGF**β**1, TGF**β**2 and PDGFAB induces morphological changes in DFT cells

TGFβ1, TGFβ2 and PDGF ligands demonstrated potent effects on DFT proliferation and migration; characteristics associated with mesenchymal pathway activation in cancer cells. Cancer cells with a mesenchymal phenotype are known to adopt alternative cellular morphologies (35). As cells progress through the EMT spectrum, they become flattened, elongated, and amorphous. We performed immunocytochemistry on DFT cell lines treated with TGFβ1, TGFβ2 and PDGFAB to observe changes in morphology. Treated cells were labelled for F-actin, a key cytoskeletal protein, and images shown are representative of multiple culture experiments (Figure 5, Supplementary figure 5).

**Fig. 5.**
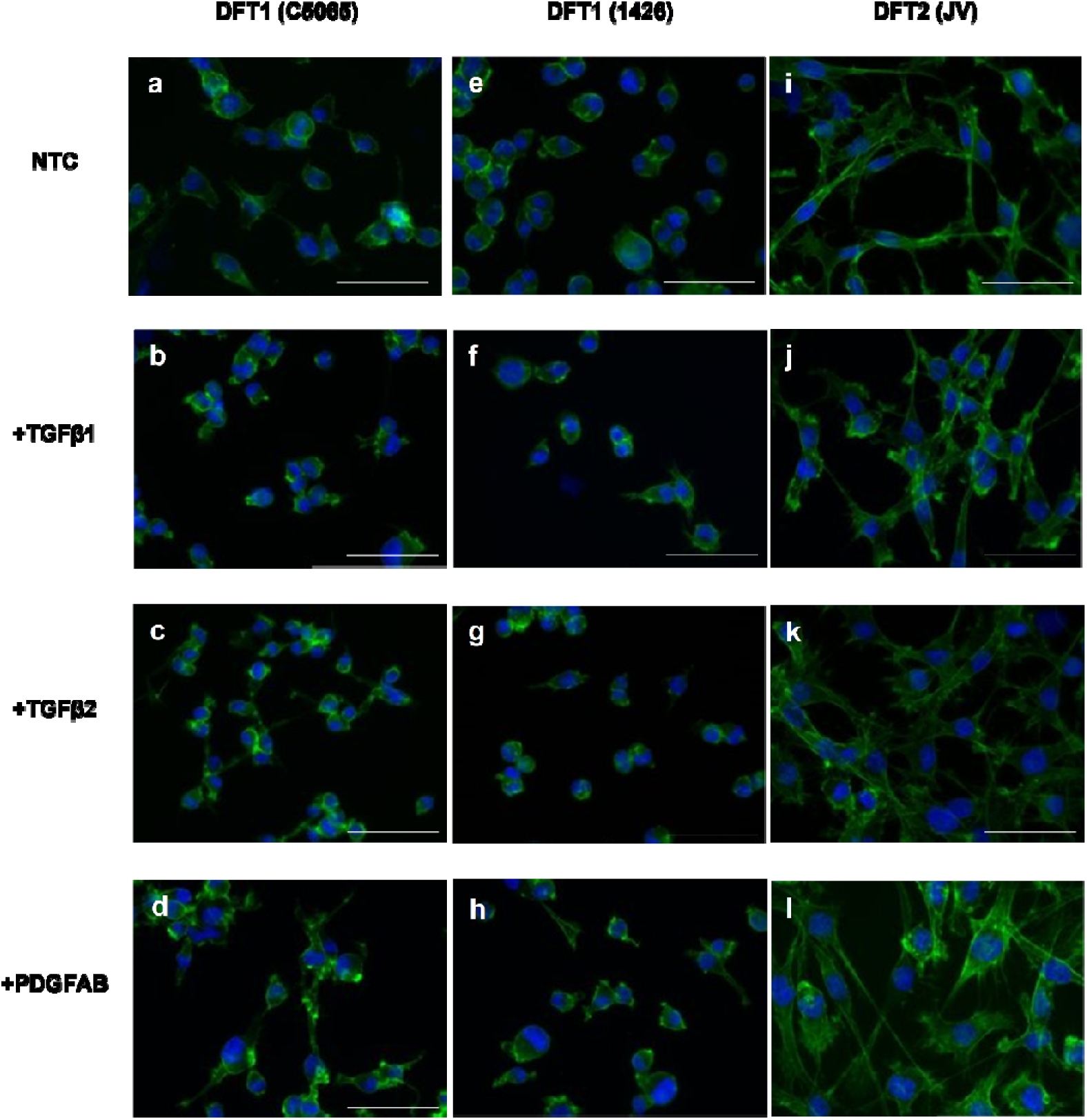
Cytoskeletal staining of treated DFT cell lines. (A-L) DFT cell line C5065 (A-D), 1426 (E-H) and DFT2 cell line JV (I-L) were treated with POIs for 48 hours and stained for actin with phalloidin-488 (green). Nuclei were coloured with 4′,6-diamidino-2-phenylindole (DAPI) (blue). No treatment controls (NTC) were included as a comparison. Stained coverslips were visualised and imaged on the BX50 fluorescent microscope at 40x magnification. The two imaged channels were later merged and colourised using Fiji (image J) analysis software (36). Scale bars represent 50 µm

DFT1 cells (C5065) treated with TGFβ1, TGFβ2 and PDGFAB for 48 hours exhibited extended cytosolic projections and elongation (Figure 5A-D). A second DFT1 cell line, 1426, also appeared elongated after treatment with TGFβ1 and TGFβ2 (Figure 5E-H). DFT2 cells treated with TGFβ1, TGFβ2 and PDGFAB exhibit a flattened, amorphous form (Figure 5I-L). Furthermore, TGFβ2-treated cells exhibited a visible rearrangement of F-actin into long, parallel fibres, extending across the length of the cell (Figure 5K). The morphological changes observed in DFT1 and DFT2 cells are characteristic of altered mesenchymal properties in cancer cells (35).

### TGF**β** pre-treatment potentially increases vulnerability to immune-mediated killing

Previous experiments demonstrated enhanced immune cell infiltration into the tumour microenvironment of previously vaccinated devils compared to naive devils (21). It is not yet known whether the mesenchymal DFT phenotype, which is characterized by elevated levels of TGFβ, PDGF, and PDGF receptors and was stimulated in these tumours, provides immune protection or heightened immunogenicity. To explore this question, we co-cultured DFT cells pretreated with TGFβ1, TGFβ2, and PDGFAB with varying ratios of devil PBMCs for 4 and 18 hours and assessed ATP production as a proxy of cell proliferation, viability and immune-mediated killing (Figure 6, A-D). Pre-treatment of DFT1 cells with TGFβ2 led to reduced ATP production in the presence of PBMCs. Similarly, increased PBMC-mediated DFT2 killing or cell cycle arrest was observed following TBFβ1 and TGFβ2 pre-treatment. These findings, along with the decrease in cell proliferation after TGFβ treatment (Figure 2B, E, H, K), suggest that TGFβ has tumour-suppressing effects on DFT cells.

**Fig. 6.**
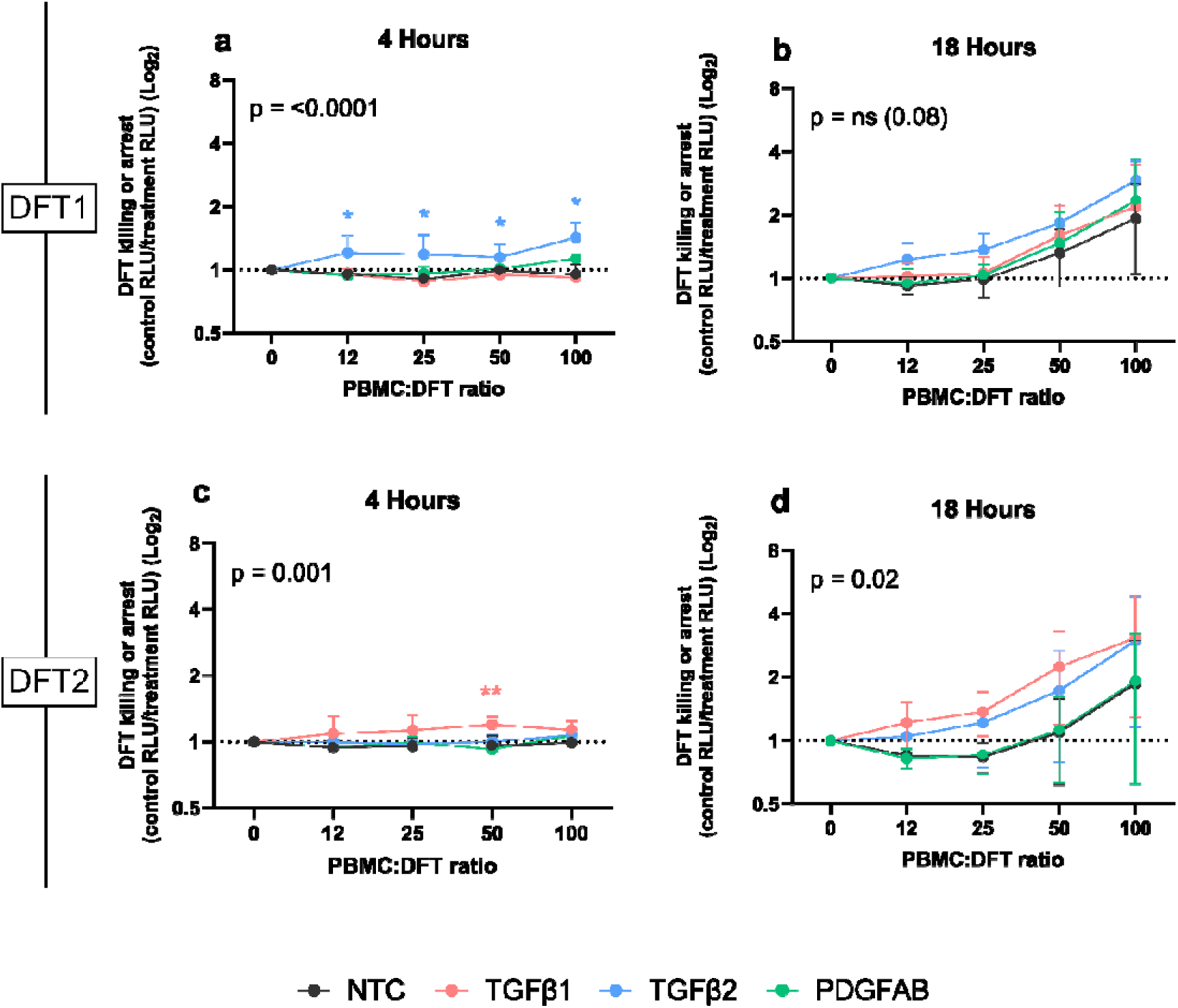
Bioluminescence imaging (BLI)-based DFT, PBMC co-culture assay. (A-D) DFT cells treated with POI’s were co-cultured with Tasmanian devil PBMC’s at increasing ratios. Bioluminescence was measured as a proxy of DFT ATP production, and therefore viability or proliferation, at 4 (A-B) and 18 (C-D) hours. Increased PBMC-mediated killing or cell arrest was estimated by the RLU in the no-PBMC control well of each treatment condition divided by the RLU in the experimental well of the same treatment condition. Each value represents the mean and standard deviation using PBMCs from three different devils for no treatment control (NTC) and treated samples. The main treatment effect compared to the no treatment control was assessed by one-way ANOVA and presented as p-values on graphs. Dunnett’s multiple comparisons test was used to compare treatment at individual cell ratios. Significance was set at p<0.05 and is indicated by asterisks: where * =p<0.05; **=p<0.01; and ***=p<0.001. Asterisks are colour co-ordinated with the respective treatment. Graphs were created using GraphPad Prism version 9.0.0

### TGFβ1 receptor inhibition suppresses DFT1 and DFT2 cell proliferation

Transforming growth factor-beta (TGFβ) is recognized for its dual role in cancer. In the early stages of tumour development, TGFβ functions as a tumour suppressor by inhibiting cell proliferation and inducing apoptosis (37). In contrast, during advanced stages, TGFβ facilitates tumour progression and metastasis through EMT (38, 39). In our study, treatment with TGFβ1 and TGFβ2 significantly inhibited proliferation in both DFT cell lines, with a more pronounced effect in DFT1 cells (Figure 2B, E, H, K). This response led us to investigate the mechanistic aspects of TGFβ signalling by blocking the TGFβ1 receptor with Galunisertib, a pharmaceutical inhibitor of the TGFβ1 receptor. Galunisertib exhibited an IC50 of 7 μM in DFT2 cells and approximately 100 μM in DFT1 cells (Figure 7A-B). The effect of Galunisertib on DFT1 cells was negligible, while TGFβ1 receptor inhibition strongly suppressed DFT2 proliferation (Figure 7C-D). These results suggest a contrasting role for TGFβ1 signalling in DFT1 and DFT2 cells.

**Fig. 7.**
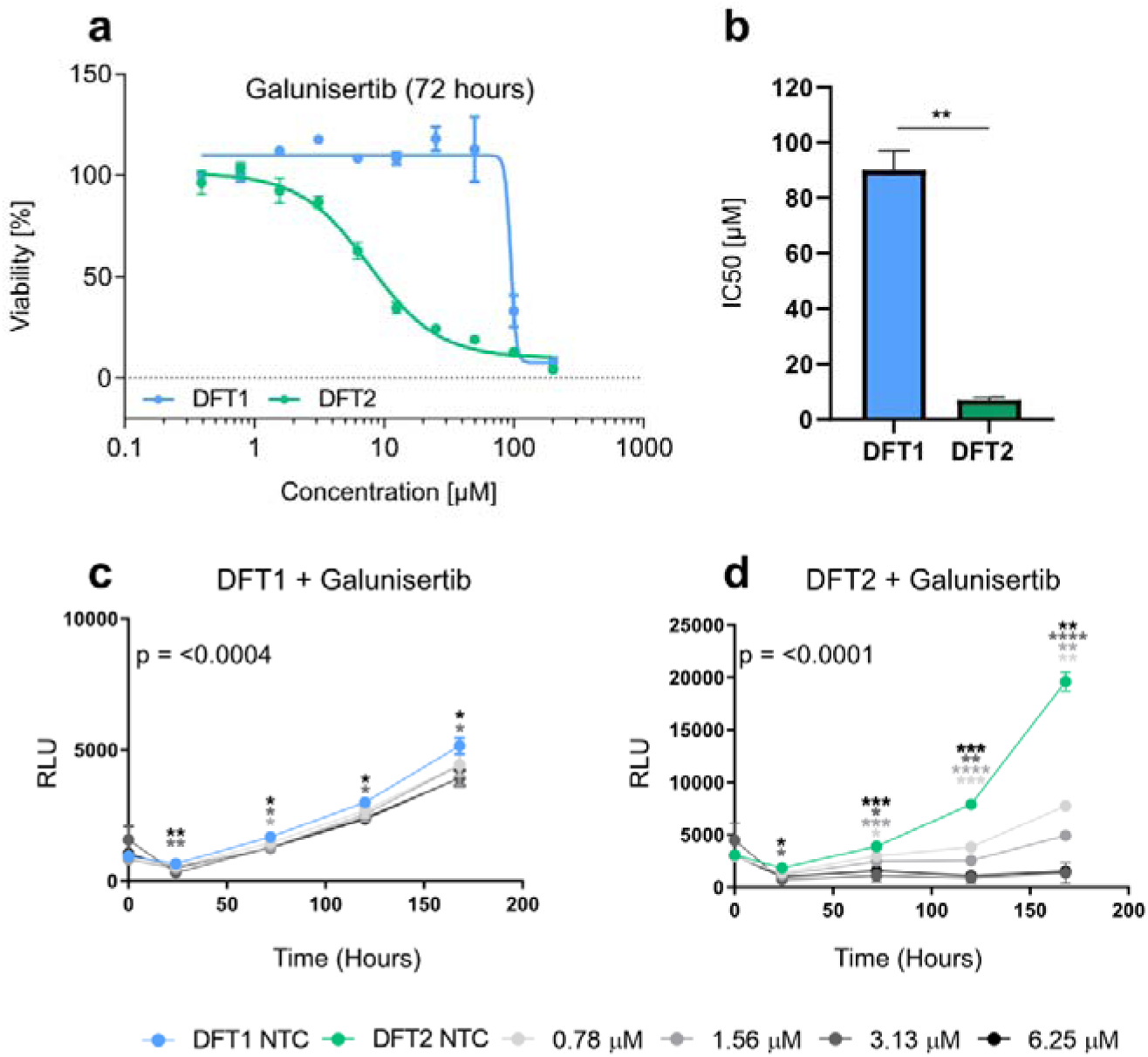
TGFβ1 receptor inhibition. (A-B) Cytotoxicity Assay: DFT1 cells and DFT2 cells were treated with serially diluted Galunisertib (TGFβ1 receptor inhibitor) for 72 hours. IC50 values were calculated through non-linear regression analysis. Statistical significance comparing IC50 values of DFT1 and DFT2 was measured using an unpaired t-test. (C-D) Bioluminescence-based proliferation assay: DFT1 cells and DFT2 cells were treated with serially diluted Galunisertib for 168 hours. Relative luminescence was measured every 48 hours as a proxy of cell number. Data for each treatment concentration were plotted as relative light units (RLU) over time. Statistical significance compared to the no treatment control (NTC; 0 ng/mL) was measured using a two-way ANOVA with Dunnett’s multiple comparisons test. P-values on graphs represent the significance of the overall interaction effect between time and treatment concentration. Statistical significance was set at p<0.05 and is indicated by asterisks where * =P<0.05, **=P<0.01, ***=P<0.001 and ****=P<0.0001. Asterisks are colour co-ordinated with the respective treatment. Each point represents the mean of data in triplicate with error bars representing the standard deviation. Graphs were created using GraphPad Prism version 9.0.0

## Discussion

Previous transcriptomic studies of DFT biopsies have demonstrated mesenchymal plasticity following prophylactic vaccination and changes in the immune environment (21). These phenotypic changes were associated with immune evasion, prompting studies to investigate how phenotypic plasticity affects DFT oncogenesis. Here, we sought to understand the drivers of mesenchymal plasticity in DFT cells by treating cells in culture with NRG1, IL16, PDGFAA, PDGFAB, TGFβ1 and TGFβ2; signalling proteins previously associated with varied DFT phenotypes (9, 21, 31). To measure plasticity in the treated cells, we considered increased migratory capacity, change in proliferative capacity and altered mesenchymal cellular morphologies as indicators of mesenchymal plasticity (40, 41). All the signalling proteins tested significantly affected at least one of these functions in DFT cells, highlighting a crucial role of the tumour microenvironment in controlling DFT phenotype in Tasmanian devils.

It was previously hypothesised that the inherent plasticity of DFT cancers in response to immune factors is partially attributed to their Schwann cell origin (21). Supporting this idea, the changes to treated DFT cells observed in this study were similar to those described in the literature regarding Schwann cell plasticity (42–44). Under homeostatic conditions, Schwann cells are associated with peripheral axons and support the conduction of action potentials through myelin production and maintenance. During peripheral nerve injury or degeneration, Schwann cells dissociate from the axon and transition into a repair Schwann cell that is functionally suited to promoting axon repair through a variety of mechanisms (45). Activation of this repair Schwann cell program involves similar pathways to well-characterised EMT programs, and has been described as partial EMT (46). Both processes are driven and sustained by signalling factors in the microenvironment (47, 48).

NRG1 and IL16 are factors that were previously associated with myelinating (semi-differentiated) phenotype of DFT cells (21). Our findings show that treatment with both proteins enhanced the migratory capacity of both DFT1 and DFT2 but had no effect on proliferation. Indeed, in other species, IL16 has been shown to increase cell migration via its secondary receptor CD9, which is expressed at high levels on DFT cells (21, 49–51). Comparatively, NRG1 exists in multiple isoforms with pleiotropic roles in Schwann cells that vary from maintenance of myelin protein expression, to directing the long-term survival and migration of repair Schwann cells during peripheral nerve repair (52–54). Thus, the net effect of NRG1 treatment on DFT cells may depend on the specific NRG1 isoform used. Indeed, genes associated with signalling through ERBB3, a receptor activated by NRG1, are upregulated in DFT1 and this signalling pathway acts as a driver of DFT1 proliferation (21, 31). We did not observe increased proliferation in response to the NRG1 isoform used in this study (ENSSHAT00000019687.2, extracellular domain).

Other signalling molecules analysed in this study are associated with mesenchymal (de-differentiated) phenotypes in DFT1 and DFT2 cells. PDGFAA and PDGFAB are homo- and heterodimers capable of activating PDGF receptors. They are known drivers of EMT in multiple human cancers and have important roles in the Schwann cell repair program (22, 43). Two studies by Stammnitz *et al*. (2018, 2023) discovered copy number amplification of *PDGFRA* in DFT2 tumours and *PDGFRB* in DFT1 tumours, suggesting a critical role of these pathways in DFT oncogenesis (6, 9). Accordingly, we found that PDGFAA/AB increased the proliferation and migration of both cancers. In the context of peripheral nerve repair, Schwann cells rapidly proliferate to replace damaged ones, which differs from the classical definitions of EMT and aligns with the repair Schwann cell program (55, 56). This suggests that PDGFAB could be activating conserved Schwann cell pathways in DFT cells to promote growth and migration.

TGFβ drives EMT in many human cancers (38). In the early stages of tumour development, TGFβ functions as a tumour suppressor by inhibiting cell proliferation and inducing apoptosis (37). Likewise, proliferation of DFT1, which exhibits a more epithelial cell-like phenotype, was inhibited by TGFβ1. This decreased proliferation coincided with an increased vulnerability *in vitro* to PBMC-mediated DFT killing or cell cycle arrest that could have been initiated by rapid innate lymphoid cells, such as NK cells (57). During later stages of tumour development, TGFβ facilitates tumour progression and metastasis through EMT (38, 39). Similarly, DFT2 was inhibited by a suppressor of TGFβ1 receptor signalling, indicating a pro-proliferative role for TGFβ1 in this less differentiated cancer.

In other species, mesenchymal activation of cancer cells confers immune evasive properties, and therapeutic mesenchymal stimulation has been explored as an avenue for preventing graft rejection in humans following transplantation (58). Specifically, mesenchymal cells exhibit resistance to killing by CD8^+^ T cells and NK cells through multiple mechanisms including upregulation of immune checkpoint molecules (23, 59). TGFβ in the tumour microenvironment can also inhibit NK cell cytotoxic activity by suppressing metabolic pathways and activating receptors (57, 60). The dual role of TGFβ in cancer, suppressing tumour growth while also dampening immune responses, emphasizes its complexity as a pleiotropic cytokine. The observed production of TGFβ in DFTs suggests a mechanism by which these tumours evade immune detection and destruction, independent of MHCI regulation. DFT tumours may have developed mechanisms to counteract inhibition of proliferation by autologous TGFβ production, or alternative ligands, through upregulation of PDGFR signalling pathways. This adaptive strategy underscores the complexity of immune evasion in DFT cancers and demonstrates the importance of understanding these mechanisms for developing tumour interventions.

Transmissible DFT cells must exhibit highly evolved immune evasion strategies to persist across genetically diverse hosts and, further, to persist despite vaccination (19). Here, we have demonstrated that DFT cancers are capable of phenotypic plasticity in response to cytokines present in the tumour microenvironment. In devils, tumour phenotypes are likely the summative effect of multiple, interacting pathways, providing challenges in replicating these phenotypes in an *in vitro* environment. Nonetheless, previous transcriptional analyses and the functional data provided in this study suggest a mechanism by which changes in the DFT microenvironment induced by anti-tumour responses drives phenotypic plasticity, allowing tumour cells to evade immune defences (21). This phenotypic plasticity reveals DFT1 and DFT2 as moving targets for anti-tumour immunity.

## Statements and Declarations

The authors wish to thank Anne-Maree Pearse (DPIPWE), Kate Swift (DPIPWE), Alex Kreiss (UTAS) and Ruth Pye (UTAS) for the DFT cell lines used in this study. We thank Jocelyn Darby and Kelsie Raspin for technical support for the experiments presented in this manuscript, and Greg Woods, Bruce Lyons and Aaron Smith for their helpful comments.

## Author Contributions

All authors were involved in the study’s conception and design. KM and AS conducted the experiments, AF developed the cell lines, and KM, AS, and AP performed data collection and analysis. KM and AS wrote the initial draft of the manuscript, with all authors providing feedback on draft versions. All authors read and approved the final manuscript.

## Data Availability

All data supporting the figures in this manuscript is provided or is available by email request.

## List of common abbreviations

DFTD: Devil Facial Tumour Disease DFT1 Devil Facial Tumour 1
DFT2: Devil Facial Tumour 2
EMT: Epithelial – Mesenchymal Transition
TME: Tumour Microenvironment
NK: Natural Killer cell
MHCI/II: Major Histocompatibility Complex 1 and 2 IL Interleukin
TGFβ: Transforming Growth Factor Beta
TGFβ1/2: Transforming Growth Factor Beta 1/2
TGFBR1/2: Transforming Growth Factor Beta Receptor 1 and 2
PDGFA/B: Platelet Derived Growth Factor A and B
PDGFRA/B: Platelet Derived Growth Factor Receptor A and B
NRG1: Neuregulin 1
STAT1/3: Signal Transducer and Activator of Transcription 1 and 3

