## Supplementary figure for "Differentially expressed growth factors and cytokines drive phenotypic changes in transmissible cancers"

Cellular and Molecular Life Sciences

Kathryn G. Maskell*^1^, Anna Schönbichler*^2^, Andrew S. Flies^1^, Amanda L. Patchett^1,3^

^1^Menzies Institute for Medical Research, University of Tasmania, Tasmania, Australia.

^2^Institute for Animal Breeding and Genetics, University of Veterinary Medicine Vienna, Vienna, Austria.

^3^Agriculture and Food, Commonwealth Scientific and Industrial Research Organisation (CSIRO), Tasmania, Australia.

*Authors contributed equally to the work

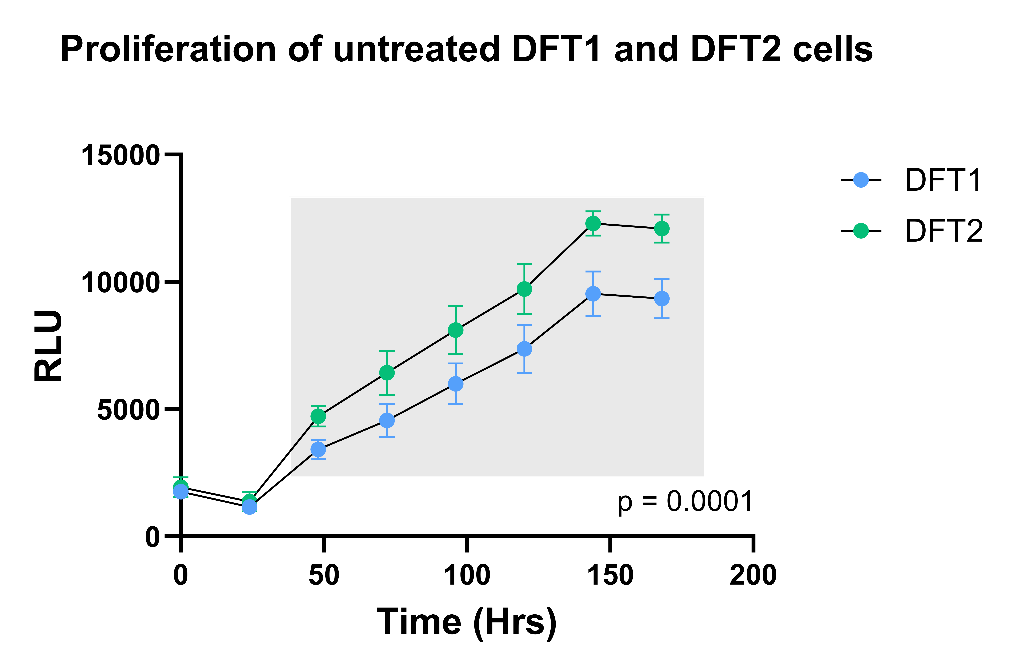

**Sup. Fig. 1** Proliferation rates of untreated DFT1 and DFT2 cells stably transfected with luciferase. Untreated DFT1 cells and DFT2 cells were plated in triplicate. Relative light units (RLUs) were measured every 24 hours as a proxy of cell number. Data for each cell line were plotted as relative light units (RLU) over time (hours). Statistical significance was measured using a two-way ANOVA with Sidak’s multiple comparisons test. The P-value on the graph represents the significance of the overall interaction effect between time and cell line. Statistical significance of multiple comparisons was set at p<0.05 and is indicated on the graph by the grey box, within which the comparison between DFT1 and DFT2 was significant (p<0.0001). Each point represents the mean of data in quintuplicate with error bars representing the standard deviation. Graph was created using GraphPad Prism

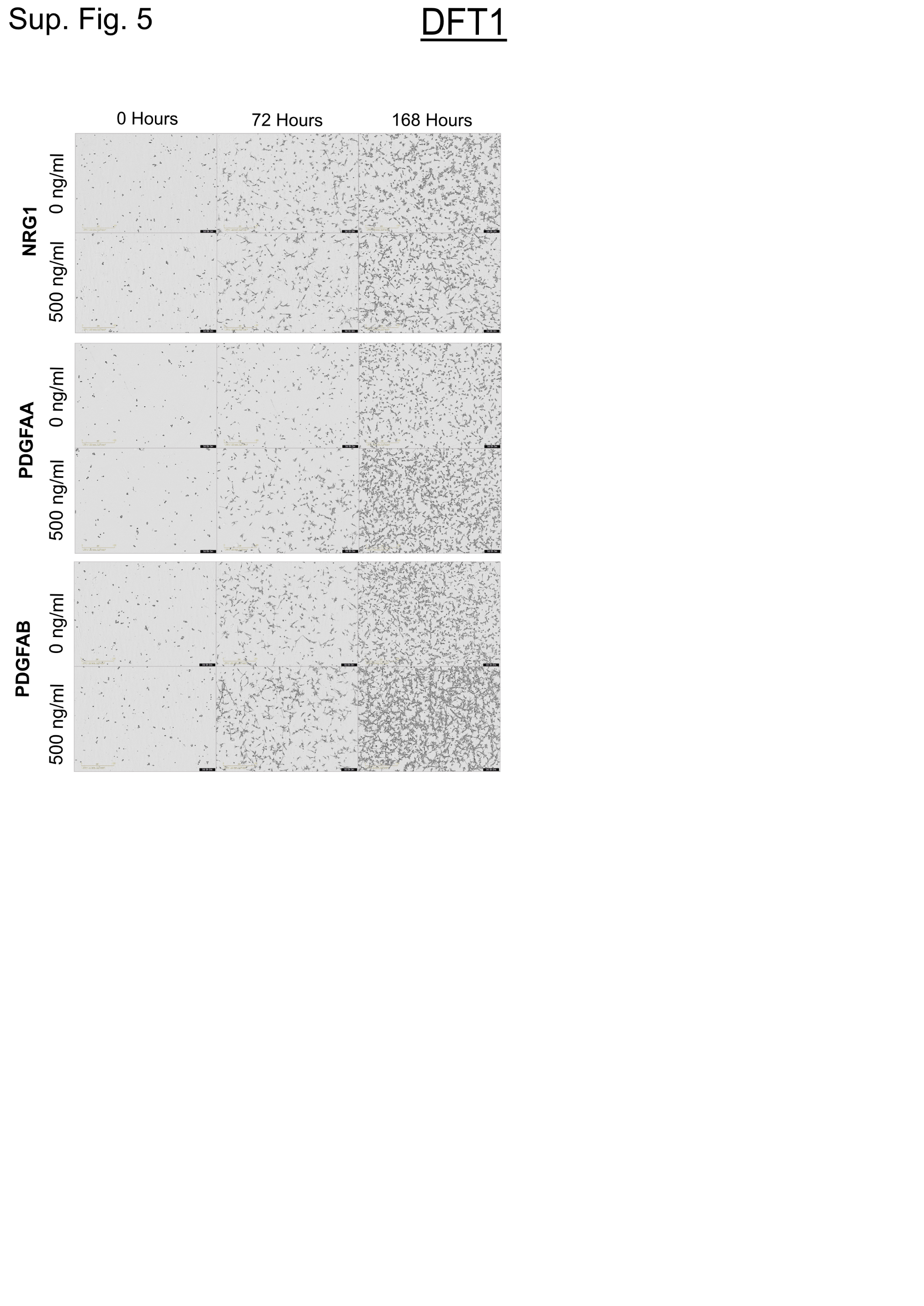

**Sup. Fig. 2** High concentration proliferation assay images of DFT1 cells treated with POIs for 168 hours. DFT1 cells were treated with (A) NRG1, (B) PDGFAA and (C) PDGFAB for 168 hours. Cell density and abundance was visualized by scanning the assay plates in an *Incucyte* Live-Cell Analysis system after seeding (0 hours), 72 hours and 168 hours of treatment. Picture displaying cells in the lowest concentration (0 ng/ml) and highest concentration (500 ng/ml) are shown

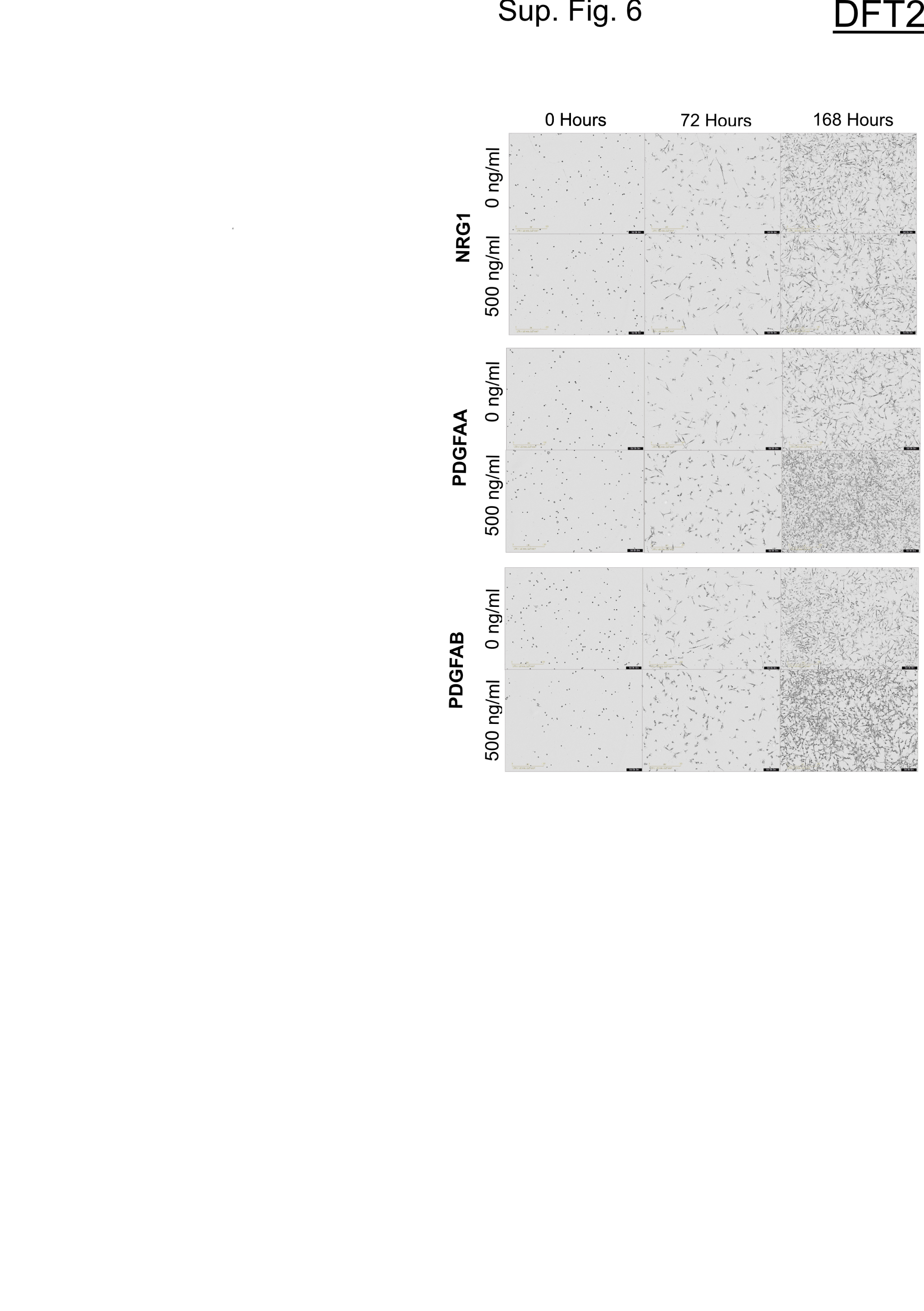

**Sup. Fig 3** High concentration proliferation assay images of DFT2 cells treated with POIs for 168 hours**.** DFT2 cells were treated with (A) NRG1, (B) PDGFAA and (C)PDGFAB for 168 hours. Cell density and abundance was visualized by scanning the assay plates in an *Incucyte* Live-Cell Analysis system after seeding (0 hours), 72 hours and 168 hours of treatment. Picture displaying cells in the lowest concentration (0 ng/ml) and highest concentration (500 ng/ml) are shown

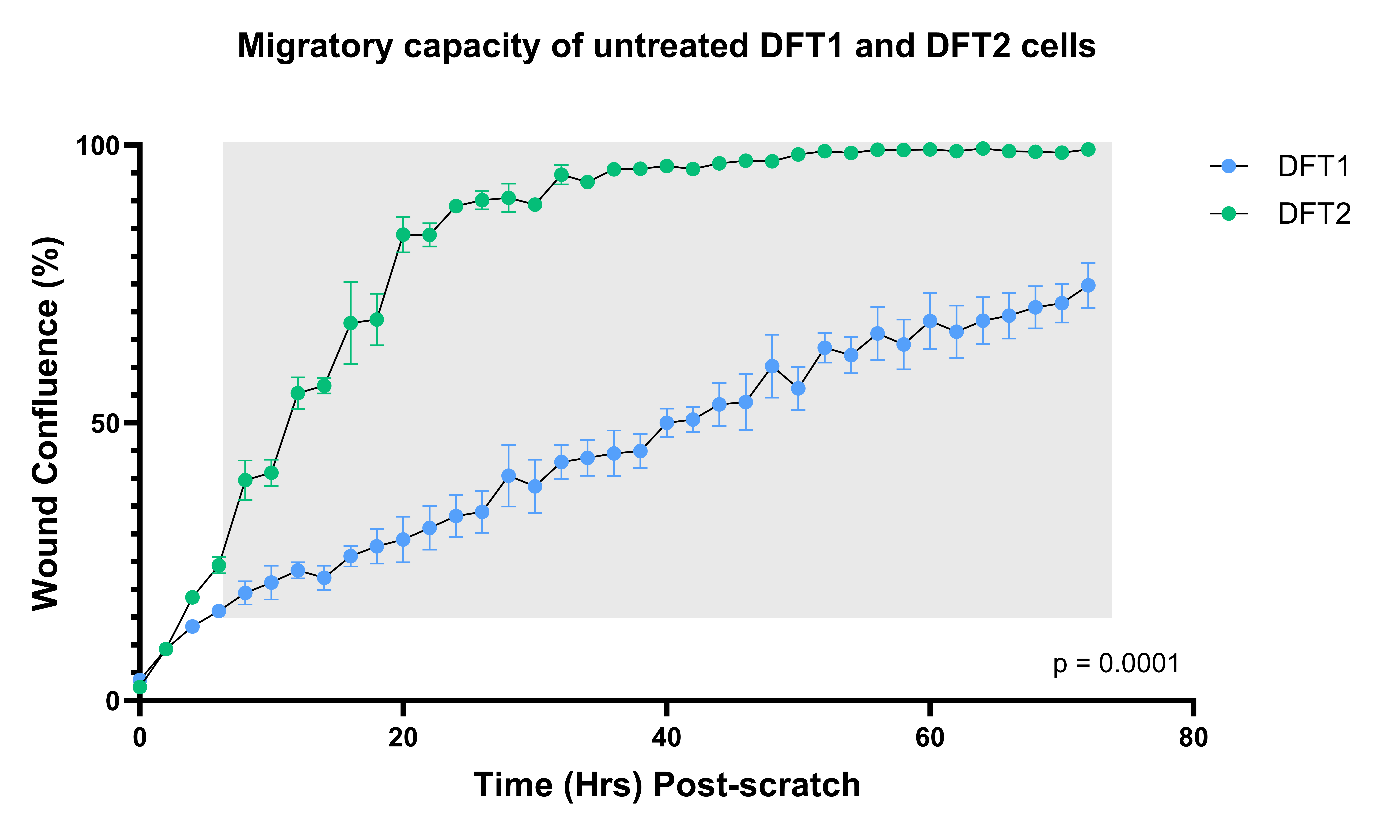
**Sup. Fig. 4** Scratch plate migration assay. Untreated DFT1 and DFT2 cells were plated in triplicate on 0.3 mg/mL Matrigel. A 700-800 μm scratch wound was created down the centre of the well and media added. Images were taken every 2 hours. Each point represents mean wound confluency (%) with error bars showing standard deviation. Statistical significance was tested by a two-way ANOVA followed by Sidaks multiple comparisons test with significance set to p<0.05. The P-value for the interaction effect is given on the lower right-hand side of the graph, and multiple comparisons tests were significant (p<0.001) at all time points within the grey box. Graph was created using GraphPad Prism

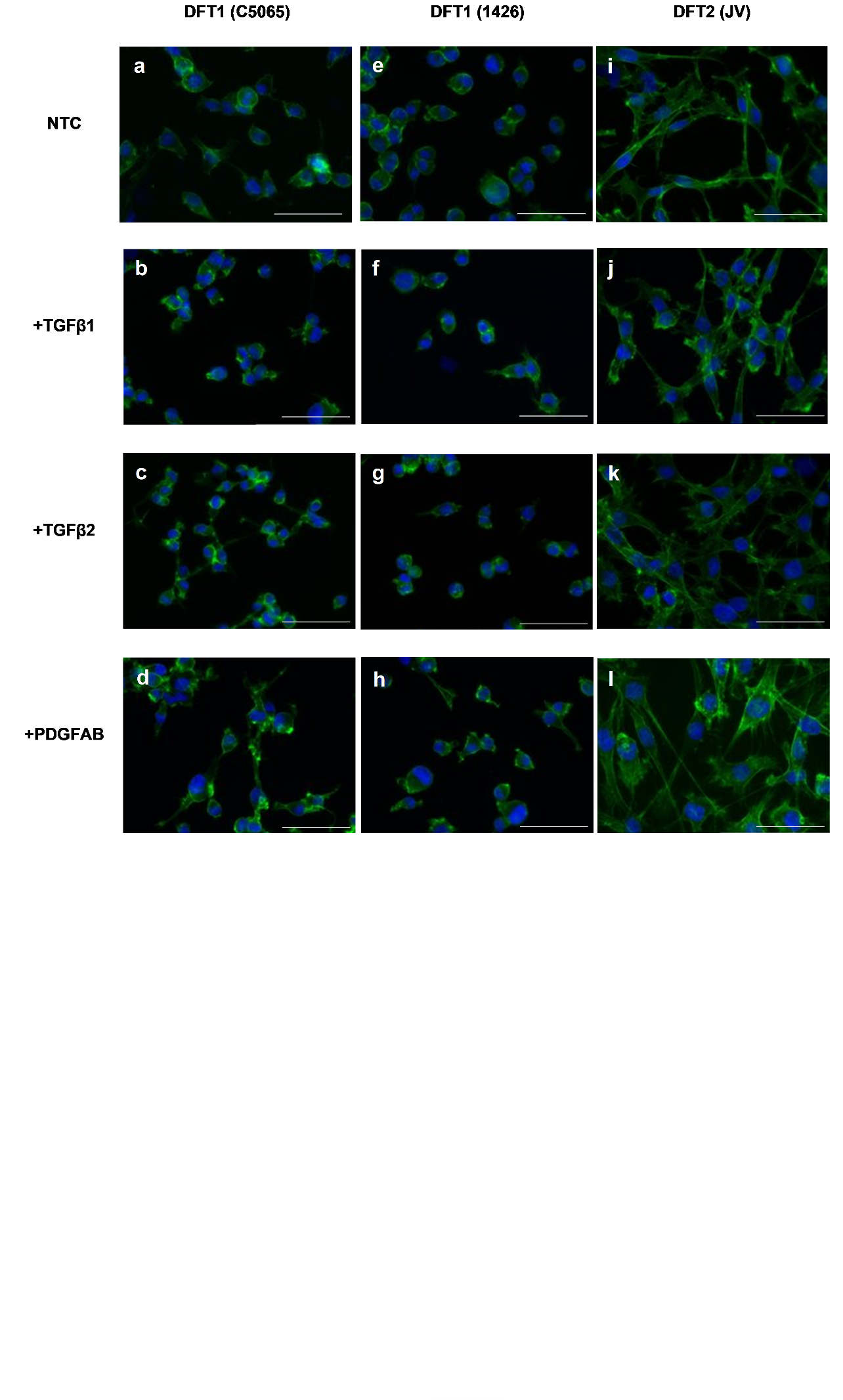

**Sup. Fig. 5** Cytoskeletal staining of DFT cell lines. (A-L) DFT cell line C5065 (A-D), 1426 (E-H) and DFT2 cell line JV (I-L) were treated with POIs for 24 hours and stained for actin with phalloidin-488 (green). Nuclei were coloured with 4′,6-diamidino-2-phenylindole (DAPI) (blue). at No treatment (NTC) controls were included as a comparison. Stained coverslips were visualised and imaged on the BX50 fluorescent microscope at 40x magnification. The two imaged channels were later merged and colourised using Fiji (image J) analysis software (Schindelin et al., 2012). Scale bars represent 50 µm
